# Predictive modeling of susceptibility to substance abuse, mortality and drug-drug interactions in opioid patients

**DOI:** 10.1101/506451

**Authors:** Ramya Vunikili, Benjamin S. Glicksberg, Kipp W Johnson, Joel Dudley, Lakshminarayanan Subramanian, Khader Shameer

## Abstract

Opioid addiction causes high degree of morbidity and mortality. Preemptive identification of patients at risk of opioid dependence and developing intelligent clinical decisions to deprescribe opioids to the vulnerable patient population may help in reducing the burden. Identifying patients susceptible to mortality due to opioid-induced side effects and understanding the landscape of drug-drug interaction pairs aggravating opioid usage are significant, yet, unexplored research questions. In this study, we present a collection of predictive models to identify patients at risk of opioid abuse, mortality and drug-drug interactions in the context of opioid usage. Using publicly available dataset from MIMIC-III, we developed predictive models (opioid abuse models a=Logistic Regression; b=Extreme Gradient Boosting and mortality model= Extreme Gradient Boosting) and identified potential drug-drug interaction patterns. To enable the translational value of our work, the predictive model and all associated software code is provided. This repository could be used to build clinical decision aids and thus improve the optimization of prescription rates for vulnerable population.

## Introduction

Drug overdose is the leading cause of accidental deaths in the US with 52,404 lethal drug overdoses in 2015. Opioid addiction is driving this epidemic with 20,101 overdose deaths related to prescription pain relievers and 12,990 overdose deaths related to heroin in 2015. The overdose death rate in 2008 was nearly four times that in 1999 and the sales of prescription pain relievers in 2010 were four times those in 1999 ([1]-[2]). Also, a study done by Jeffery et al., ([6]) highlights the fact that despite all the increased attention to opioid abuse and awareness of risks, the opioid use and average daily dose have not substantially decreased from the peaks. Drug overdose continues to be an alarming public health problem and thus, it needs immediate attention. However, a part of this problem could be addressed if we can precociously identify those subjects who are the most susceptible to adverse events when given opioids. We provide a solution to this by using simple yet robust machine learning techniques involving classification algorithms. In addition to this, we’ve explored the interactions between opioids and other drugs that could result in increased incidence of a particular side effect. In order to discover the relation between the interactions and the incidence of side effects we’ve performed K-Means clustering. As aptly described in Khader et al., ([10]), this study combines the robustness of both statistical analysis and machine learning techniques. It also exemplifies the utility of publicly available biomedical datasets and its application for improving public health as emphasized by Khader et al., ([9]).

Even after being acknowledged as one of the major issues, opioid epidemic remains eclipsed from the artificial intelligence communities in healthcare. Che et al., ([4]) is one of the very few attempts done to classify subjects based on opioid usage. This study categorizes subjects into three groups (short term, long term and opioid dependent users) based on the number of prescriptions given. Here, opioid dependent users refer to those who are diagnosed with “opioid dependence”. This study describes two classification tasks: a) whether a short term user will turn into a long term user and b) whether a long term user is an opioid dependent user. One issue with such a type of classification is that the study is ignoring the possibility of a short term user developing the symptoms of opioid dependence. When a subject is prescribed opioids only a few times but with high dosages the subject could still be prone to adverse effects. Another point to be noted in this study is that the best performing model for identifying opioid dependent users is a deep learning model that uses Recurrent Neural Network (RNN). As highlighted by Dudley et al., ([8]), it is a well-established fact that deep learning models need to be trained on large datasets for better performances. However, as the number of subjects who experienced opioid dependence symptoms in Che et al., ([4]) was only 749, this study has randomly generated 14 datasets by downsampling non-opioid-dependent subjects which formed two-thirds of the dataset and then trained the RNN model. This might not be the most technically robust way to generate data. Even with such a random generation the accuracy of the model is found to be 76.07% with a recall of only 52.05%. That means, the chances of identifying a long term subject who could be prone to opioid dependence using this model is better than tossing a fair coin by a mere margin of 2%.

Also, as Dudley et al., ([8]) pointed out, deep learning models are often regarded as models lacking interpretability in healthcare. To overcome all these issues, our study advocates the use of traditional machine learning models to achieve better classification accuracies by extracting data in a more robust way.

## Materials and Methods

### Dataset

The MIMIC-III dataset consists of details of 46,520 subjects at the Beth Israel Deaconess Medical Center, Boston, Massachusetts. Among these, 29,959 subjects were identified with prescriptions of opioids such as Morphine, Meperidine, Codeine, Buprenorphine, Hydromorphone, Methadone, Fentanyl, Oxycodone, Oxymorphone and Hydrocodone. Further, 1,405 subjects out of these were prescribed Naloxone, which is an anti-narcotic medication known for its usage as opioid overdose reversal drug. In a few cases, Buprenorphine could also be prescribed in combination with Naloxone to minimize the possibility of opioid dependence.

### Cohort Selection

All the subjects with opioid prescriptions were divided into 8 age groups. Age of the subjects was calculated based on their date of birth and the date of prescription issued. In case of multiple prescriptions issued for a subject, the latest prescription was considered for determining his/her age. The statistics of each of these age groups is presented in Table 1.

**Table 1.** Statistics of subjects in different age groups.

| Age Group | Age Range (Years) | Total No. Of Subjects | No. Of Subjects With Side Effects |
| --- | --- | --- | --- |
| 1 | <13 | 269 | 0 |
| 2 | 13 - 19 | 253 | 7 |
| 3 | 20 - 40 | 2949 | 254 |
| 4 | 41 - 50 | 3273 | 203 |
| 5 | 51 - 65 | 8507 | 251 |
| 6 | 66 - 75 | 5974 | 26 |
| 7 | 76 - 85 | 5906 | 7 |
| 8 | >85 | 2861 | 1 |

In order to identify subjects with side effects, we’ve checked the diagnoses reports of every subject prescribed with opioids for symptoms related to overdose and/or dependence using the International Classification of Diseases, Ninth Revision (ICD 9) codes. A few of the ICD 9 codes and categories are listed in Table 2. A total of only 749 subjects were identified to have side effects.

**Table 2.**
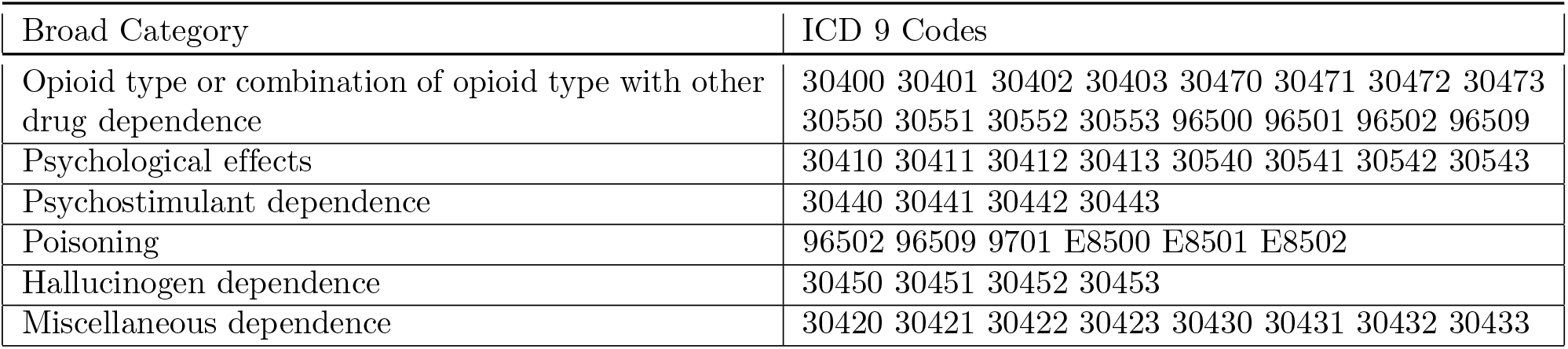
List of ICD 9 codes used for identifying subjects with adverse events.

| Broad Category | ICD 9 Codes |
| --- | --- |
| Opioid type or combination of opioid type with other drug dependence | 30400 30401 30402 30403 30470 30471 30472 30473 30550 30551 30552 30553 96500 96501 96502 96509 |
| Psychological effects | 30410 30411 30412 30413 30540 30541 30542 30543 |
| Psychostimulant dependence | 30440 30441 30442 30443 |
| Poisoning | 96502 96509 9701 E8500 E8501 E8502 |
| Hallucinogen dependence | 30450 30451 30452 30453 |
| Miscellaneous dependence | 30420 30421 30422 30423 30430 30431 30432 30433 |

## Data Extraction Methodologies

### Feature Selection

A total of 26 features were chosen to represent the selected cohort. The target variable, *SIDE_EFFECTS_FLAG*, is set to 1 if the subject is diagnosed with any of the adverse events listed in Table 2 and 0 otherwise. The gender of a subject is represented by a binary variable - 0 for female and 1 for male. For subjects with one or more Naloxone prescriptions, the *ANTI_NARCOTIC* flag is set to 1 and for those with prescriptions of all other opioids under study, the *NARCOTIC* flag is set to 1. Every opioid is allocated a discrete variable to represent the total number of prescriptions of that particular opioid given to each subject. In addition, the total number of anti-narcotic (Naloxone) and narcotic (opioids excluding Naloxone) prescriptions are also represented by two discrete variables. If a subject has stayed in Intensive Care unit (ICU) then the binary flag, ICU, is set to 1 and 0 otherwise. The mortality status of the subject is indicated by another binary variable, *EXPIRE_FLAG*. It is triggered if the subject is no more. Also, the age group of every subject is represented using one-hot encoding. Finally, feature normalization was done by performing an affine transformation on each feature so that all the values in the dataset are in the range of [0,1]. Figure 1 shows the correlation of features. It can be observed that the target variable, *SIDE_FFECTS_FLAG*, has the highest positive correlation with *TOTAL_ANTLNARCOTIC_PRESCRIPTIONS* and *ANTL_NARCOTIC* flag. Intuitively, this makes sense because a subject would be treated with anti-narcotics when adverse events start to show up. Also, among opioids, the number of prescriptions associated with *BUPRENORPHINE* and *METHADONE* have a relatively higher positive correlation with the target variable. Similarly, the *EXPIRE_FLAG* is observed to have the highest positive correlation with *MORPHINE*.

**Figure 1.**
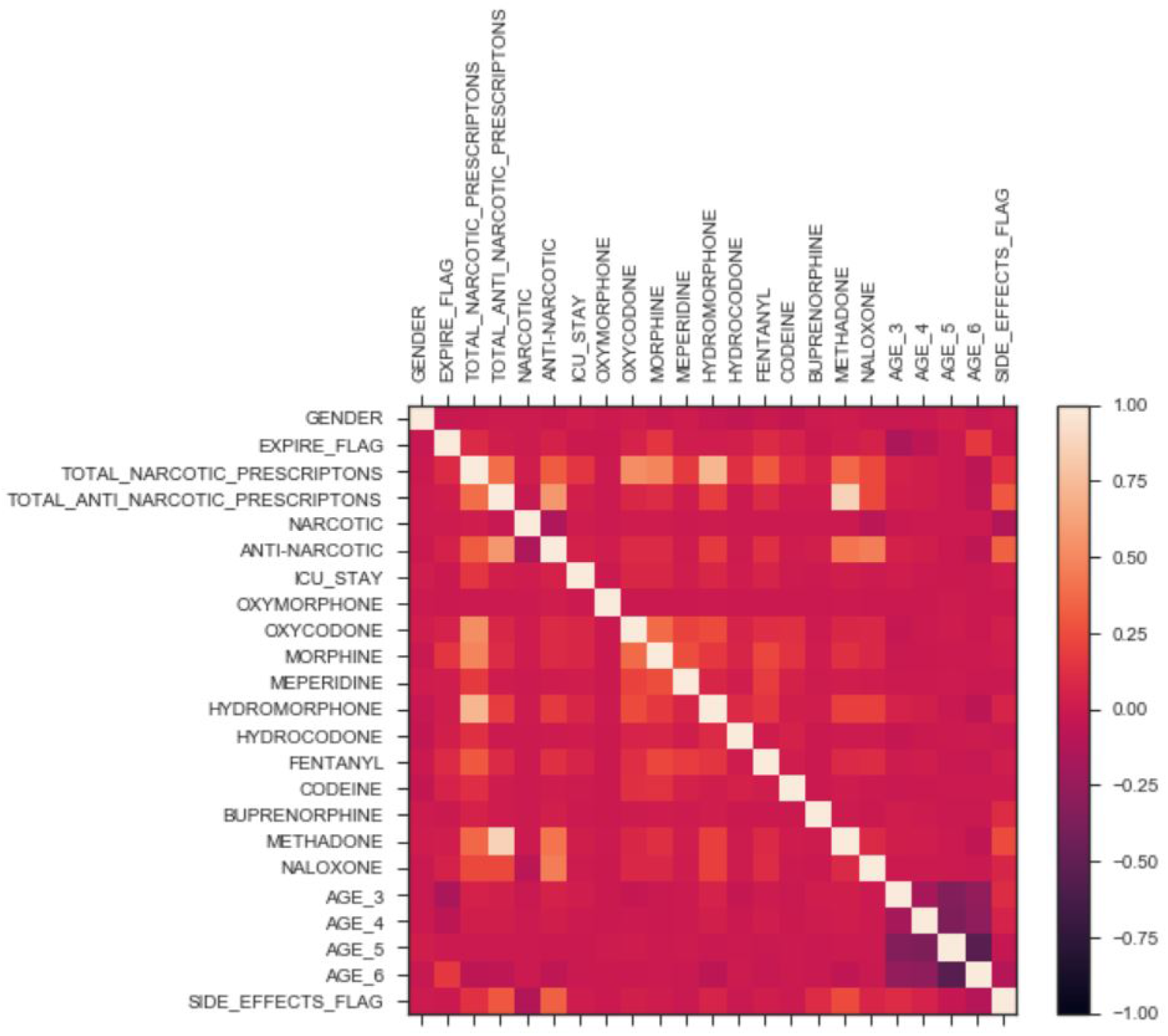
Correlation of features.

### Dealing with class imbalance

It can be observed from Table 1 that there is a huge imbalance between subjects with side effects and those with no side effects. Running a classification algorithm on such a data would result in overfitting the model and hence it will learn to predict the majority class. As a result, the classification accuracy might be high even when the number of true positives for subjects with triggered *SIDE_EFFECTS_FLAG* is terribly low. We’ve taken two steps to address this problem.

a. **Downsampling majority class** Among 749 subjects identified with side effects, only 15 belonged to age groups 1, 2, 7 and 8. On the other hand, these age groups accounted for, approximately, 10,000 samples of majority class. Although excluding these age groups has resulted in a much better ratio of subjects with side effects to those with no side effects (734:19969 vs 749:29959), the data is still highly imbalanced.
b. **SMOTE - Oversampling minority class:** In order to deal with the high class imbalance in the data, Synthetic Minority Oversampling Technique (SMOTE) developed by Chawla et al., ([3]) was used. This algorithm works by choosing the nearest neighbors of data with minority class label and upsamples them. This method was used after performing Linear Discriminant Analysis (LDA) on the data which provided evidence that both the classes were quite separable from each other. Implementing this algorithm not only led to the expansion of the dataset in a statistically robust way but also minimized the imbalance in the dataset.

### Addressing the issue of sparse features

As described earlier, quite a number of features were based on the opioids given to the subjects. A few opioids like Morphine were prescribed very often while the other opioids such as Oxymorphone were rarely prescribed. As every subject had features related to every opioid, the less frequently prescribed opioids led to sparse features. In order to have a better subset of features we’ve performed Principal Component Analysis (PCA). From Figure 2, it can be observed that the maximum variance is retained from component 6 onwards. But, the regression resulted in maximum accuracy with 11 components. Hence, the number of features have been reduced to 11.

**Figure 2.**
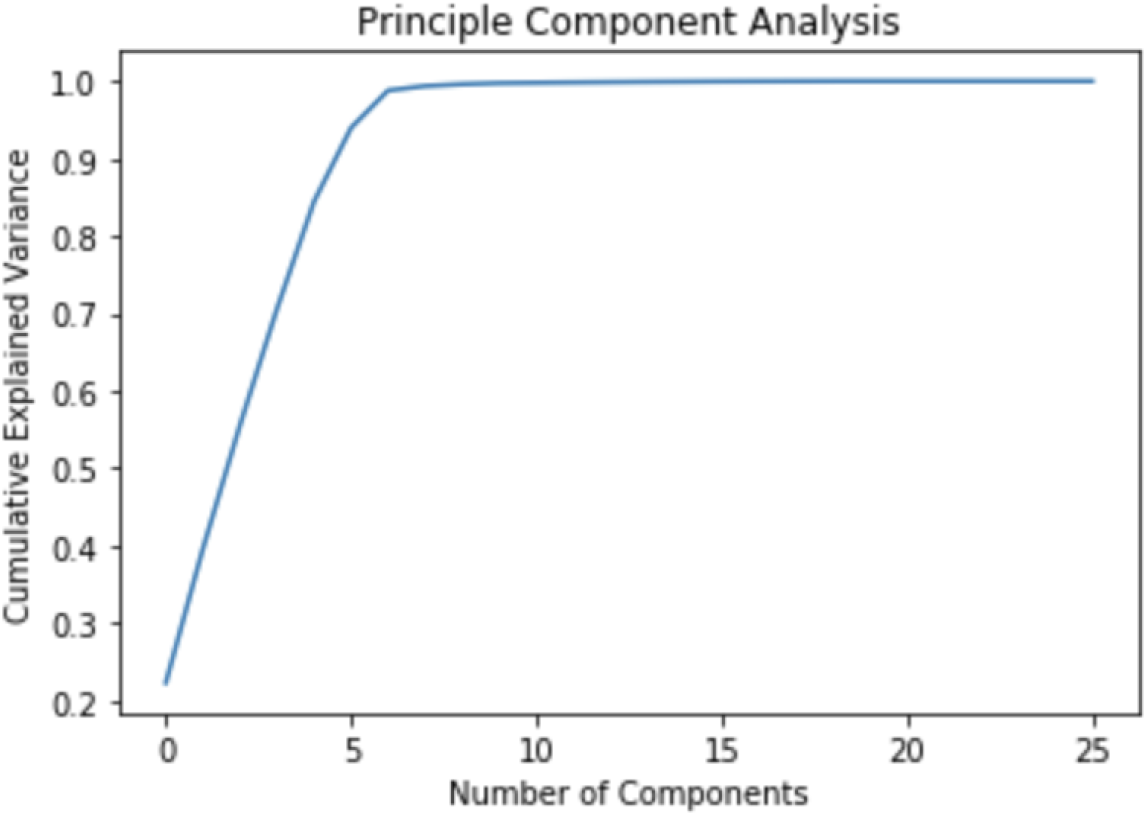
Cumulative explained variance across different principle components.

## Modelling

The entire dataset was split into 80% training set and 20% test set for running the classification models. We’ve chosen Logistic Regression with L2 regularization as a baseline and Extreme Gradient Boosting (XGBoost) developed by Chen et al., ([5]) as an enhanced model. For both the models, we have performed 10-fold cross validation on the dataset.

### Baseline model - Logistic Regression

Logistic Regression model with L2 penalty of 0.001 was run on the dataset before and after performing SMOTE and PCA. The mean AUC of 10 fold cross validation can be observed in Figure 3.

**Figure 3.**
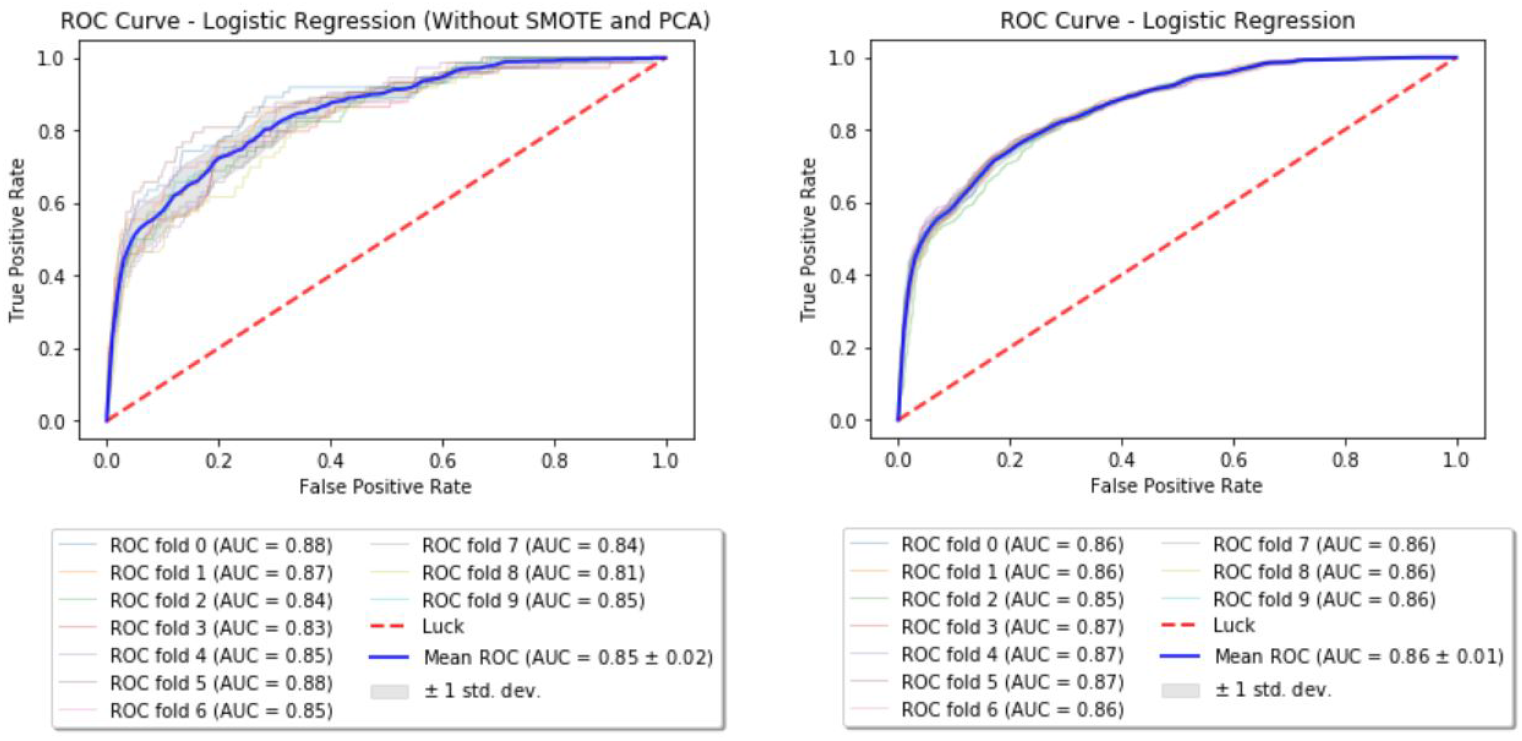
Baseline: ROC curve before after performing SMOTE and PCA.

### Enhanced model - XGBoost

For XGBoost, 20% of the training set was set aside as the validation set. Grid search was done over this validation set to get the best parameters for the model. The obtained parameters are listed in Table 3. The Receiver Operating Characteristic (ROC) curve before and after performing SMOTE & PCA for XGBoost are shown in the Figure 4.

**Figure 4.**
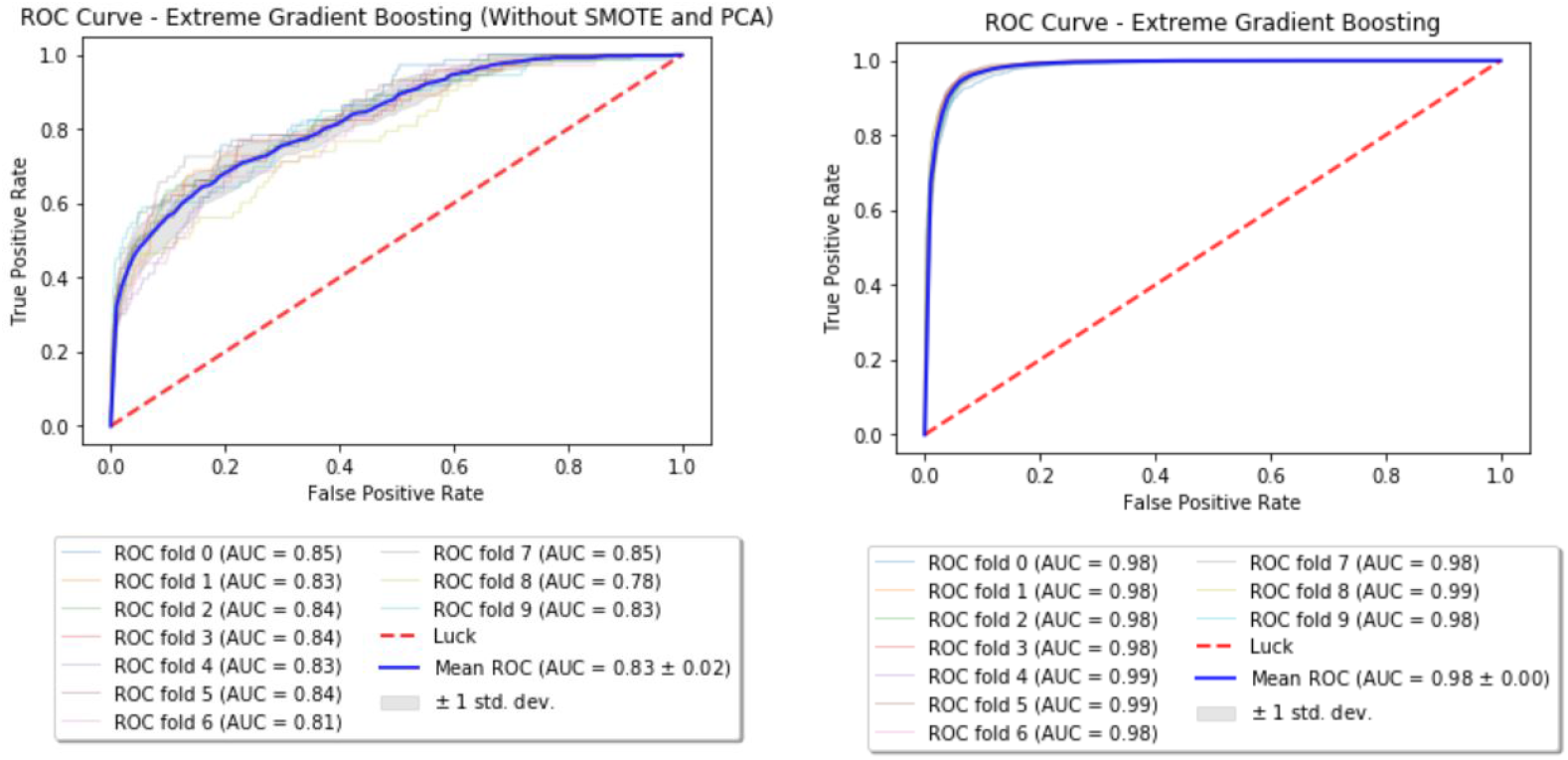
Enhanced model: ROC curve before after performing SMOTE and PCA.

**Table 3.** Summary of best parameters for XGBoost.

| Parameter | Value |
| --- | --- |
| Learning rate | 0.1 |
| Maximum depth | 8 |
| N estimators | 200 |
| Objective | binary:logistic |

### Modeling without mortality flag

As it is practical and obvious that we would like to use this model to assess patients who are alive, it makes more sense to drop the *EXPIRE_FLAG* feature and look at the performance of the model. By keeping all the data extraction and processing methodologies same, the mean AUC with XGBoost has been found to be the same as that of the XGBoost model with the *EXPIRE_FLAG*. This can be observed from Figure 5.

**Figure 5.**
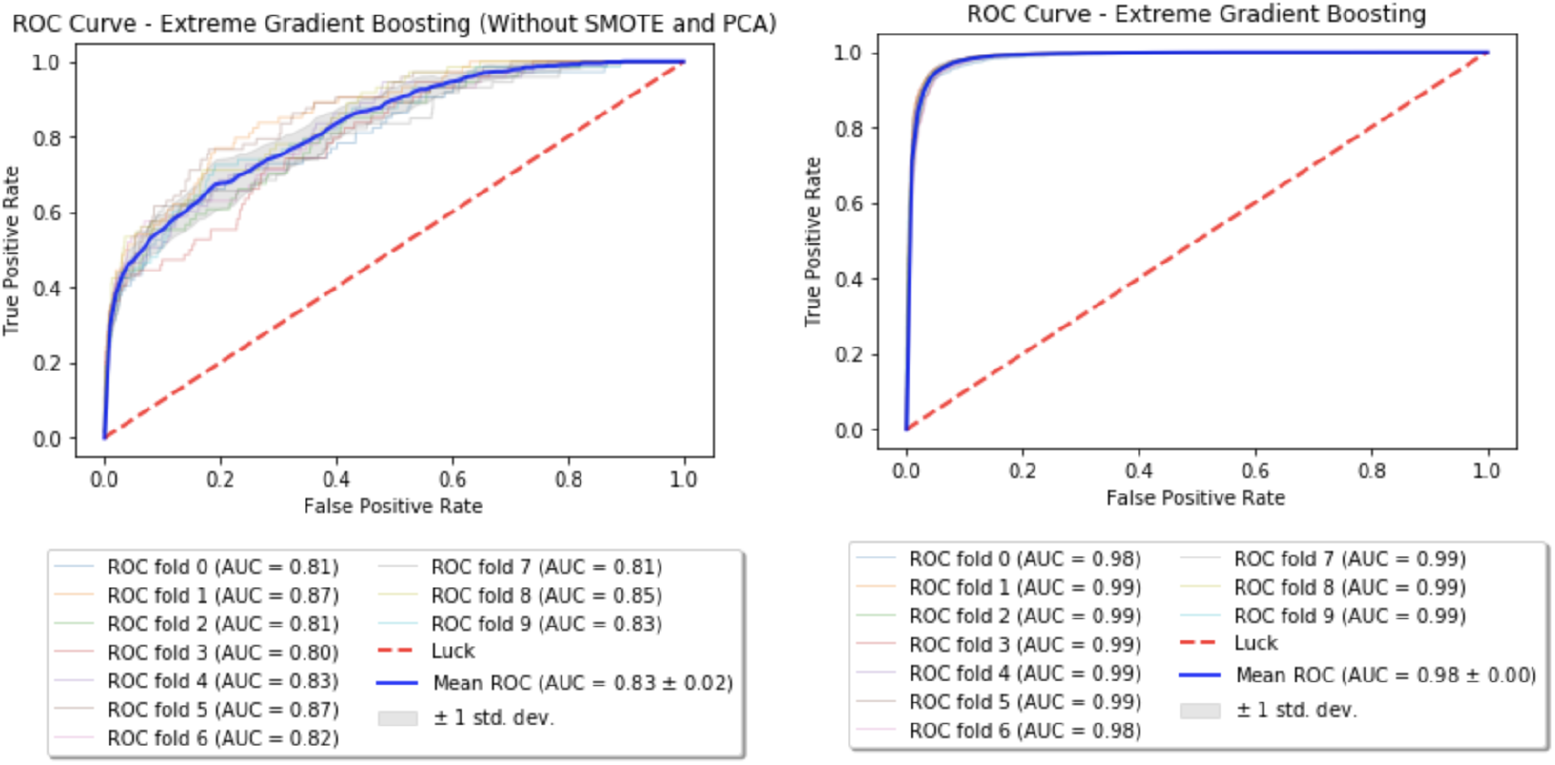
XGBoost: ROC curve before after performing SMOTE and PCA.

### Mortality as the target variable

Until now we’ve tried predicting if a subject will show side effects when prescribed opioids. But a far more fatal consequence associated with opioids is death. Being able to segregate subjects with high risk of mortality could be a huge problem in itself. Hence, to facilitate such a precocious identification we’ve run a classification algorithm on the cohort that has experienced side effects. XGBoost model was trained on 80% of these subjects (n=587) and tested on the remaining 20% (n=147). The accuracy of the model is given in the Table 4.

**Table 4.** Summary of performance.

| Model | Target Variable | Accuracy | Precision | Recall |
| --- | --- | --- | --- | --- |
| Logistic Regression | Side effects | 76.31% | 77.24% | 76.43% |
| XGBoost | Side effects | 93.26% | 92.26% | 95.47% |
| XGBoost | Mortality | 68.71% | 19.23% | 71.42% |

### Interactions between opioids and other drugs

This part of the study aims at discovering the interactions between opioids and other drugs that could lead to potential side effects in subjects. In order to carry out this study, we’ve considered the cohort of 749 subjects who were diagnosed with side effects in the previous study (including all age groups). These subjects were given at least one of the 11 opioids under consideration and 3710 other drugs put together. We’ve categorized the side effects into 7 groups and the summary is provided in Supplementary Table 1. All the opioids were assigned an index between 1 to 11 and similarly the other drugs were also indexed. For every opioid and other drug combination, the number of subjects who were diagnosed with side effects in each of the above 7 groups has been kept track of. These numbers were normalized before performing K-means clustering. From the elbow plot shown in Figure 6, the number of optimal clusters were found to be 4. Apart from the variants of regular salts like potassium chloride and sodium chloride, insulin is one important drug that has been classified into the predominant cluster.

**Figure 6.**
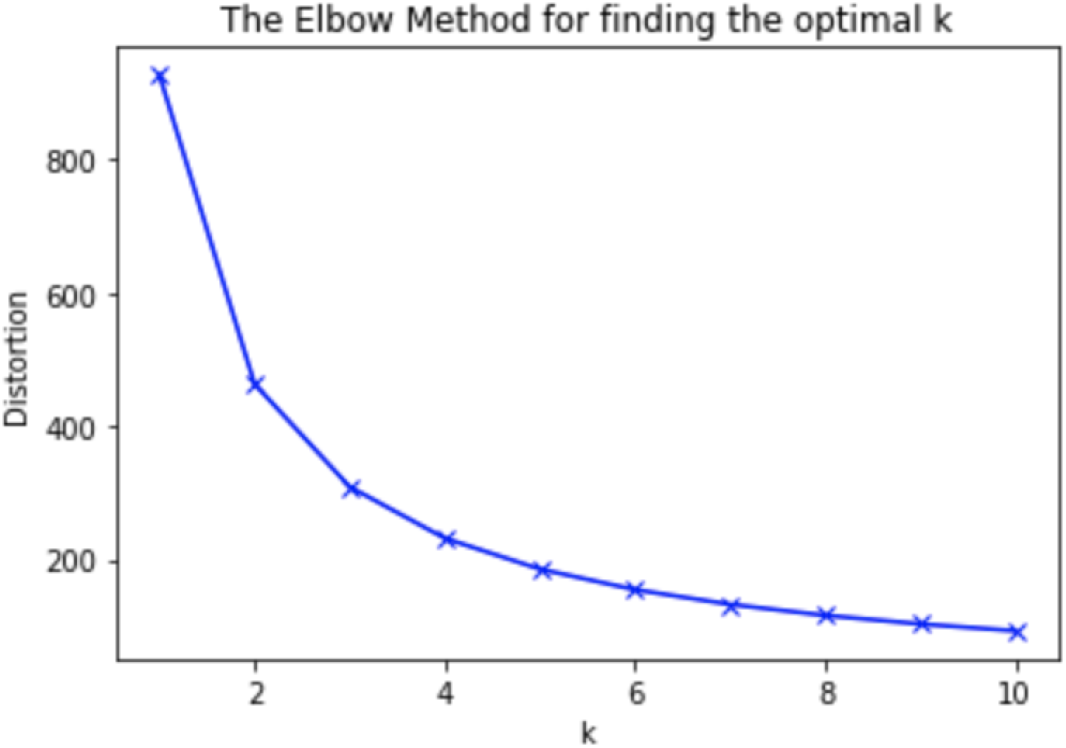
Elbow plot for K-means clustering.

## Results

The results of this study can be summarized in three sections: (a) Predictive modeling for classifying subjects susceptible to opioid abuse, (b) Predictive modeling for classifying subjects susceptible to death and (c) Interactions between opioids and other drugs.

### Predictive modeling for classifying subjects susceptible to opioid abuse

As discussed earlier, we’ve implemented two models for classifying subjects who could, possibly, be prone to adverse events upon opioid consumption. Table 4 shows that XGBoost has outperformed the Logic Regression model. This could be due to the fact every subject is associated with only a few opioids and hence only a subset of features which are related to those particular opioids are more important than the others. And since XGBoost works by sub-sampling the features, the classification accuracy of enhanced model is much higher than that of the baseline. From Figure 7, it can be observed that XGBoost has classified the number of prescriptions associated with *MEPERIDINE* as the most important feature in deciding the subject’s susceptibility to adverse events and it’s followed by *TOTAL_NARCOTIC_PRESCRIPTIONS* and *GENDER*. A more obvious result, that follows our analysis of feature correlation, is that *NALOXONE* and *MORPHINE* are also among the important contributing features.

**Figure 7.**
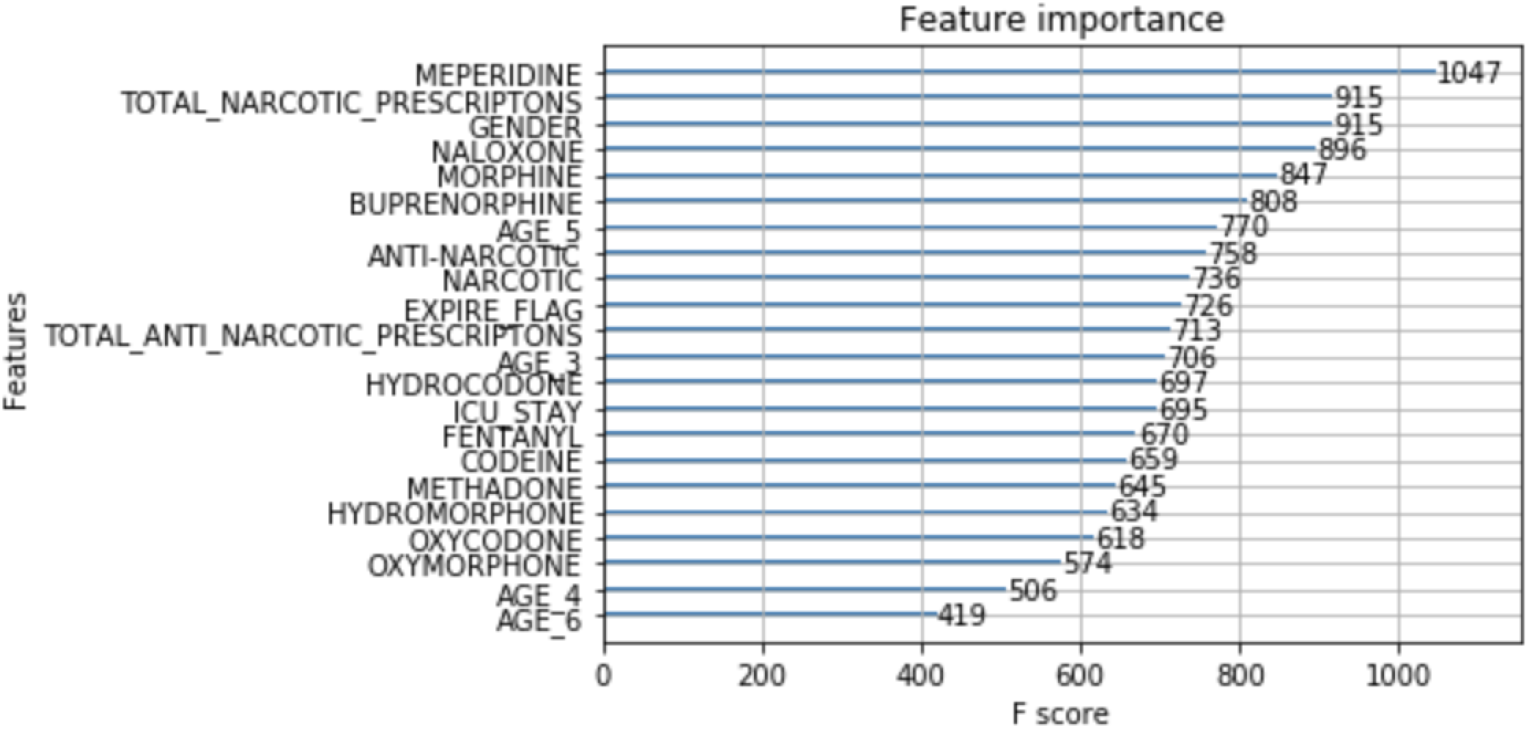
Importance of features.

Also, as desired, the XGBoost model was more sensitive in classifying subjects with adverse events than those with no adverse events. Hence, the number of true positives for label 1 are more than those for label 0 (Figure 8). In other words, the model gave a better recall score.

**Figure 8.**
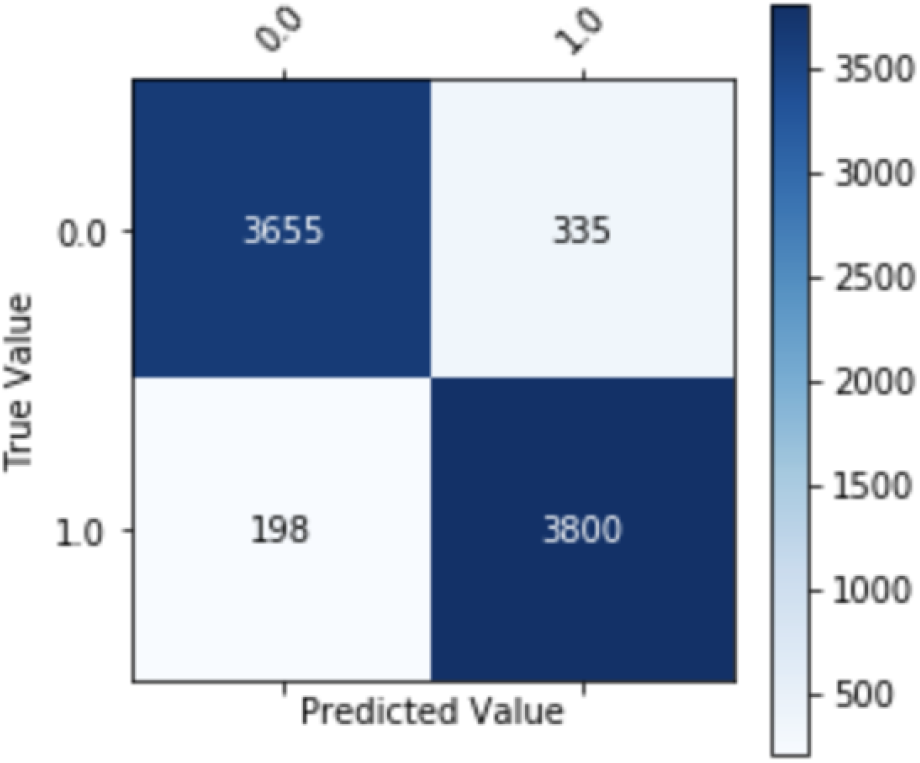
Confusion matrix.

### Predictive modeling for classifying subjects susceptible to death

Just like the previous model, it can be seen from Table 4 that the model used for classifying subjects with high risk of mortality also has a higher recall score. This implies that the model is able to classify subjects of concern with higher sensitivity.

### Interactions between opioids and other drugs

As stated earlier, insulin has been found to be in the predominant cluster associated with all categories of side effects. Not only that insulin has been used widely in subjects prescribed with opioids but also the incidence of side effects has been comparatively huge in the case of opioid and insulin combination. This observation backs up the results from two earlier studies conducted by Li et al., ([7]) and Sharma et al., ([11]) The first study states that morphine could lead to desensitization of insulin receptor signaling. This could’ve been one reason of increased usage of insulin in subjects prescribed with opioids. From the second study it can be learnt that islet cells, which are responsible for the production of insulin, do not respond in an appropriate manner to the glucose signals in subjects with opioid addiction.

## Discussion

Though the total number of subjects experiencing opioid dependence and/or adverse effects in this study is same as that in Che et al., ([4]) results show that the current models classify the subjects with a better accuracy and recall by just using traditional machine learning models. Also, our enhanced model (93.26%) has outperformed the RNN model (76.07%) in Che et al., ([4]) and can classify all subjects irrespective of the number of prescriptions given to them.

### Limitations of the current study

Having said all the above, like with any other study, there are a few drawbacks associated with this study as well. The model for predicting the mortality, unlike those for predicting the side effects, might not be robust. This is due to the fact that the reason for death of the subject remains undisclosed. Though the subject has experienced side effects, his/her death might not, necessarily, be related to opioids. This analysis of mortality prediction should be considered as a preliminary one. Further, the study of interactions between opioids and other drugs is based solely on the frequency of prescription and the frequency of incidence of side effects. As we wanted to study the correlation between the incidence of side effects and the prescription opioids/drugs, irrespective of a subject’s characteristics, we didn’t include other interactions such as protein-protein, drug-target protein etc. like that in the study done by Marnik et al., ([12]).

## Conclusion

Opioids are a class of drugs that are known for their use as pain relievers. They bind themselves to opioid receptors on nerve cells in the brain and the nervous system to mitigate pain. Addiction is one of the chronic and primary adverse events of prolonged usage of opioids. They may also cause psychological disorders, muscle pain, depression, anxiety attacks, etc. This study is intended to assist and double check the decisions taken regarding the prescription of opioids. It aims at building a predictive model to classify the subjects of interest into two categories based on their susceptibility to opioid abuse. We’ve trained two classification models, Logistic Regression with L2 regularization (baseline) and Extreme Gradient Boosting (enhanced model), to achieve this task. Results suggest that the enhanced model provides a promising approach to identify subjects who are most vulnerable to adverse events when given opioids. If employed as a reassurance technique, this study could be of tremendous help to medical practitioners in designing an appropriate action plan for their subjects before prescribing them opioids and will help combat the opioid epidemic.

## Supplementary Material

The code supporting this study is present in a private repository on Github. Please contact the corresponding authors for access. Also, the supplementary material can be referred from the same repository.

